# Diverse *Streptococcus pneumoniae* strains drive a MAIT cell response through MR1-dependent and cytokine-driven pathways

**DOI:** 10.1101/158204

**Authors:** Ayako Kurioka, Bonnie van Wilgenburg, Reza Rezaei Javan, Ryan Hoyle, Andries J. van Tonder, Caroline L. Harrold, Tianqi Leng, Lauren J. Howson, Dawn Shepherd, Vincenzo Cerundolo, Angela B. Brueggemann, Paul Klenerman

## Abstract

Mucosal Associated Invariant T (MAIT) cells represent an innate T cell population of emerging significance. These abundant cells can recognize ligands generated by microbes utilizing the riboflavin synthesis pathway, presented via the major histocompatibility complex (MHC) class I-related molecule MR1 and binding of specific T cell receptors (TCR). They also possess an innate functional programme allowing microbial sensing in a cytokine-dependent, TCR-independent manner. *Streptococcus pneumoniae* is a major human pathogen that is also associated with commensal carriage, thus host control at the mucosal interface is critical. The recognition of S. *pneumoniae* strains by MAIT cells has not been defined, nor have the genomics and transcriptomics of the riboflavin operon (Rib genes). We examined the expression of Rib genes in S. *pneumoniae* at rest and in response to metabolic stress and linked this to MAIT cell activation *in vitro.* We observed robust recognition of *S. pneumoniae* strains at rest and following stress, using both TCR-dependent and TCR-independent pathways. The pathway used was highly dependent on the antigen-presenting cell, but was maintained across a wide range of clinically-relevant strains. The riboflavin operon was highly conserved across a range of 571 S. *pneumoniae* from 39 countries dating back to 1916, and different versions of the riboflavin operon were also identified in related *Streptococcus* species. These data indicate an important functional relationship between MAIT cells and S. *pneumoniae*,which may be tuned by local factors, including the metabolic state of the organism and the antigen-presenting cell that it encounters.

**Author Summary:** *Streptococcus pneumoniae* is the leading cause of bacterial pneumonia, causes invasive diseases such as meningitis and bacteraemia, and is associated with significant morbidity and mortality, particularly in children and the elderly. Here, we demonstrate that a novel T cell population called Mucosal-associated invariant T (MAIT) cells is able to respond to a diverse range of S. *pneumoniae* strains. We found that this response was dependent on the T cell receptor (which recognises metabolites of the bacterial riboflavin biosynthesis pathway), cytokines, and the type of antigen-presenting cell. A population genomics approach was also used to assess the prevalence and diversity of the genes encoding the riboflavin biosynthesis pathway among a large and diverse collection of S. *pneumoniae.* These genes were highly conserved across a range of 571 S. *pneumoniae* from 39 countries dating back to 1916, and was also present in other related *Streptococcus* species. Given the low levels of MAIT cells in neonates and MAIT cell decline in the elderly, both of whom are at the highest risk of invasive pneumococcal disease, further understanding of the functional role of MAIT cells in host defense against this major pathogen may allow novel therapeutics or vaccines to be designed.

## Introduction

*Streptococcus pneumoniae* (the ‘pneumococcus’) is the most common cause of community-acquired pneumonia and is associated with significant morbidity and mortality, especially among young children and older adults (1,2). S. *pneumoniae*also causes invasive diseases like meningitis and bacteraemia, and upper respiratory tract infections like otitis media and sinusitis, among all age groups (3). Antimicrobial-resistant strains are widespread and pose problems in the treatment of infections, which led the World Health Organisation to include S. *pneumoniae* on their recent list of priority pathogens (4). The currently available pneumococcal conjugate vaccines prompt an immune response to polysaccharide capsules (differentiated as serotypes) and they are highly effective at preventing invasive pneumococcal disease due to vaccine-serotype strains; however, current vaccines only protect against a small number of the possible serotypes which has led to increases in the rates of disease caused by non-vaccine-serotype pneumococci are problematic (5,6). Therefore, pneumococcal disease remains a serious problem and a better understanding of the host defense against S. *pneumoniae* may allow for the design of novel therapeutics or vaccines.

There is an increasing appreciation of the role played by unconventional T cells in orchestrating early cellular events in response to invading pathogens (7). Mucosal-associated invariant T (MAIT) cells are a recently described innate T cell population, which are abundant in mucosal tissues including the lung, as well as the blood and liver (8–10). These cells conventionally express a semi-invariant T cell receptor (TCR) consisting of a Va7.2-Jα33/Jα12/Jα20 chain, paired with a limited repertoire of Vβ chains (11). This TCR is able to recognize ligands presented by the conserved major histocompatibility complex (MHC)-related protein 1 (MR1) (8). MR1 binds vitamin-B based precursors derived from the riboflavin biosynthesis pathway, which is conserved across various bacteria and fungi, but does not occur in humans (10). The ligands presented by MR1 are derived from either early intermediates in riboflavin synthesis (5-A-RU) that form adducts with other small metabolites (5-OP-RU), or the direct precursors of riboflavin (e.g. ribityllumazine (RL)-6,7-diMe) (10,12,13). Human MAIT cells are also characterized by a high expression of the C-type lectin-like receptor CD161 and the interleukin (IL)-18 receptor, and are able to respond to innate cytokines even in the absence of prior activation or TCR signaling (14,15). Upon activation, these cells produce various immunomodulatory cytokines including IFNγ, TNFα, IL-17, and Granzyme B.

MAIT cells are critical for the control of bacterial infections in mice, particularly in the lungs (16–18). For instance, aerosol infection models with *Mycobacterium bovis*Bacillus Calmette-Guérin (BCG) and *Francisella tularensis* live vaccine strain (LVS) demonstrated that MAIT cells were essential for the early control of the bacterial burden in the lung (18,19). Indeed, MAIT cells are required to coordinate both innate and adaptive arms of the immune response during pulmonary infections, as early GM-CSF production from lung MAIT cells in response to *F. tularensis* was required for the differentiation of dendritic cells, without which there was a delayed recruitment of activated CD4+ T cells (20). Thus, the rapid and innate activation of MAIT cells in response to pulmonary bacteria is critical for bridging the innate and adaptive system, shaping the local adaptive immune response.

Despite their role in pulmonary infection models, it remains unclear whether MAIT cells play a role in the defense against S. *pneumoniae* infection. Our first aim was to determine whether MAIT cells can respond to S. *pneumoniae*, and if so, to elucidate the mechanism driving such activation. We found that MAIT cells responded to S. *pneumoniae* in an MR1-dependent manner in the presence of macrophages but not monocytes, and was dependent on co-stimulation provided by innate cytokines. *In vitro* experiments and RNA-sequencing experiments also suggested that temperature and riboflavin availability affected S. *pneumoniae*, which in turn affected the responsiveness of MAIT cells. Secondly, using a population-level genomics approach, we found that the riboflavin synthesis pathway is ubiquitous and highly conserved amongst S. *pneumoniae*, which may have implications in the design of immunotherapy approaches or vaccines that target MAIT cell activity. Riboflavin operon genes were also found among other non-pneumococcal *Streptococcus*species (spp.), including S. *agalactiae* (group B streptococci), which suggests that perhaps the immunological observations made here could be extended to other human-associated *Streptococcus* spp. infections.

## Results

### *S. pneumoniae* possess a highly-conserved riboflavin synthesis operon that is upregulated with heat stress

The genes encoding the riboflavin biosynthetic enzymes of *S. pneumoniae (ribD, ribE, ribA* and *ribH)* were found to be clustered together in the same orientation in a predicted 3.4 Kb operon structure (Figure 1A). The prevalence and sequence diversity of the coding regions of the riboflavin genes were investigated in a large, global and historical genome dataset of S. *pneumoniae* isolated between 1916 and 2008 from people of all ages residing in 36 different countries. 561 (98.2%) of the S. *pneumoniae* genomes contained the riboflavin operon. Nine of the ten genomes that lacked the operon were of a single multilocus sequence type (note: ST^serotype^), ST13^14/nontypable^, and the other belonged to ST695^4^ (Supplementary Table 1). All genes in the riboflavin operon were found to be highly conserved: nucleotide and amino acid sequence identity were >99% (Table 1). The dN/dS analysis revealed a higher prevalence of synonymous versus non-synonymous mutations, supporting the importance of maintaining the riboflavin operon (Table 1).

**Figure 1.**
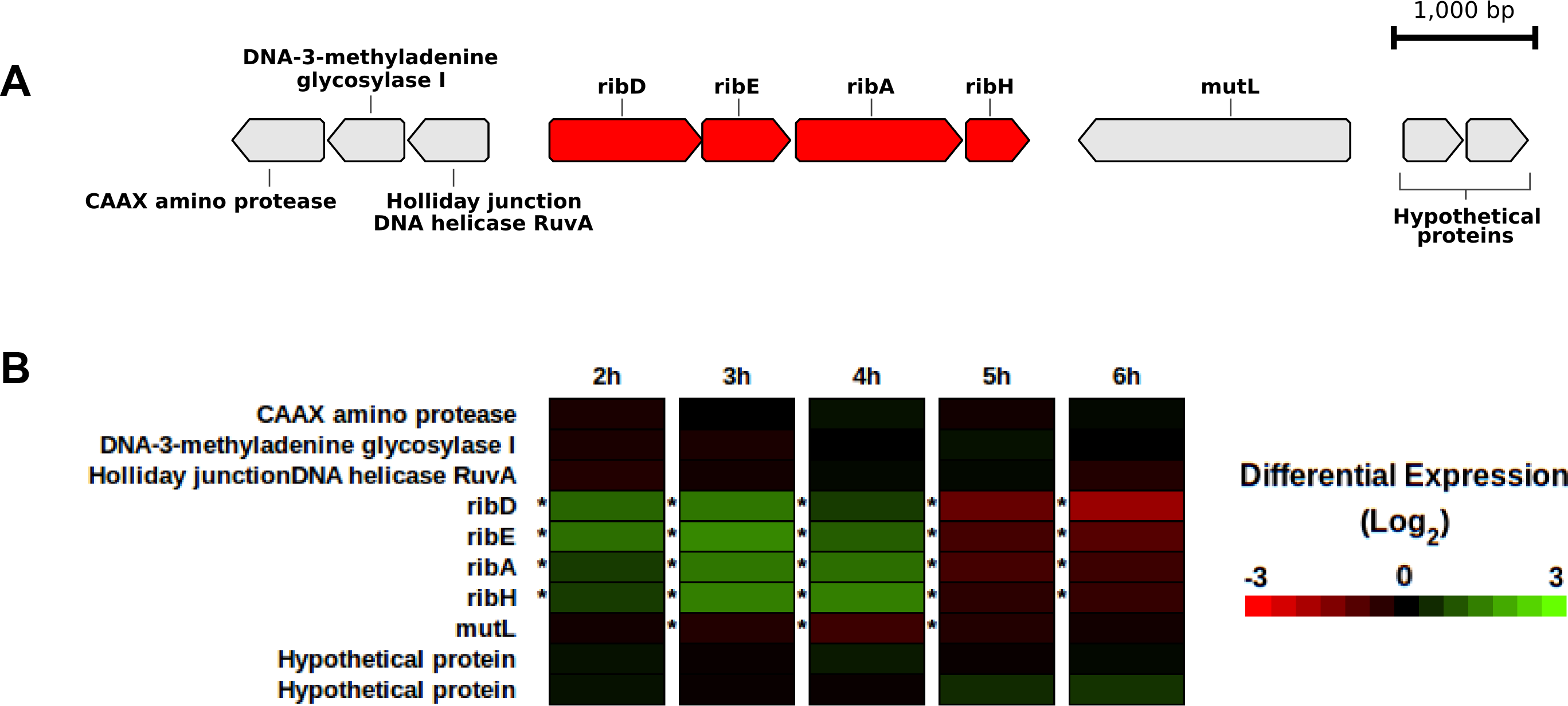
Genetic and transcriptomic data related to the riboflavin operon in S. *pneumoniae*. A) The riboflavin operon is depicted with riboflavin genes *ribD, ribE, ribA* and *ribH* (red) and flanking genes (grey). B) RNA expression data at five timepoints (2-6 hours after initial incubation) are illustrated for each riboflavin gene and the flanking genes. Genes marked with differential expression levels in green were upregulated and those in red were downregulated during 40°C incubation, as compared to normal 37°C incubation. An asterisk (*) to the left of a cell indicates a statistically significant result.

**Table 1.** Description of the four riboflavin operon gene sequences within 571 pneumococcal genomes.

| Gene | Present (%)<sup>a</sup> | Nucleotide % pairwise identity | Amino acid % pairwise identity | Mean dN/dS | Gene annotation |
| --- | --- | --- | --- | --- | --- |
| *ribD* | 559 (98.2%) | 99.6% | 99.8% | 0.36 | Diaminohydroxyphosphoribosylamino-pyrimidine deaminase |
| *ribE* | 561 (98.2%) | 99.6% | 99.9% | 0.28 | riboflavin synthase |
| *ribA* | 561 (98.2%) | 99.7% | 99.9% | 0.42 | 3,4-dihydroxy-2-butanone-4-phosphate synthase |
| *ribH* | 559 (98.2%) | 99.5% | 99.1% | 0.12 | 6,7-dimethyl-8-ribityllumazine synthase |

- a. Two genomes possessed sequence assembly gaps in *ribD* and *ribH*, and 10 genomes were missing all four of the riboflavin genes (see Results).

Sequencing was performed on RNA extracted from a pneumococcus that was incubated at a higher temperature than normal (40°C vs. 37°C). Differential expression analysis revealed that all the riboflavin operon genes were significantly up-regulated after 2-4 hours of incubation under heat stress as compared to the control (Figure 1B). Subsequently, the riboflavin operon was found to be significantly down-regulated after 5–6 hours of incubation. The concurrent increase and decrease in the expression of the four riboflavin genes suggested that these genes are transcriptionally coupled.

### MAIT cells are activated by *S. pneumoniae*

To determine whether MAIT cells were able to respond to *S. pneumoniae*, 10 Pneumococcal Molecular Epidemiological Network (PMEN) reference strains were used to probe the activation of MAIT cells in the presence of the monocytic cell line, THP1. Peripheral blood mononuclear cells (PBMCs) were cultured with paraformaldehyde (PFA)-fixed *S. pneumoniae* and THP1 cells overnight, or *Escherichia coli (E. coli)* as a positive control (Figure 2A). There was a clear production of interferon-γ (IFNγ) from MAIT cells across all strains although there was variability in the responses: 7 out of 10 strains reached sengineered to express the MAITstatistical significance (PMEN2, PMEN13, PMEN14, PMEN34, PMEN35, PMEN36, and PMEN39).

**Figure 2.**
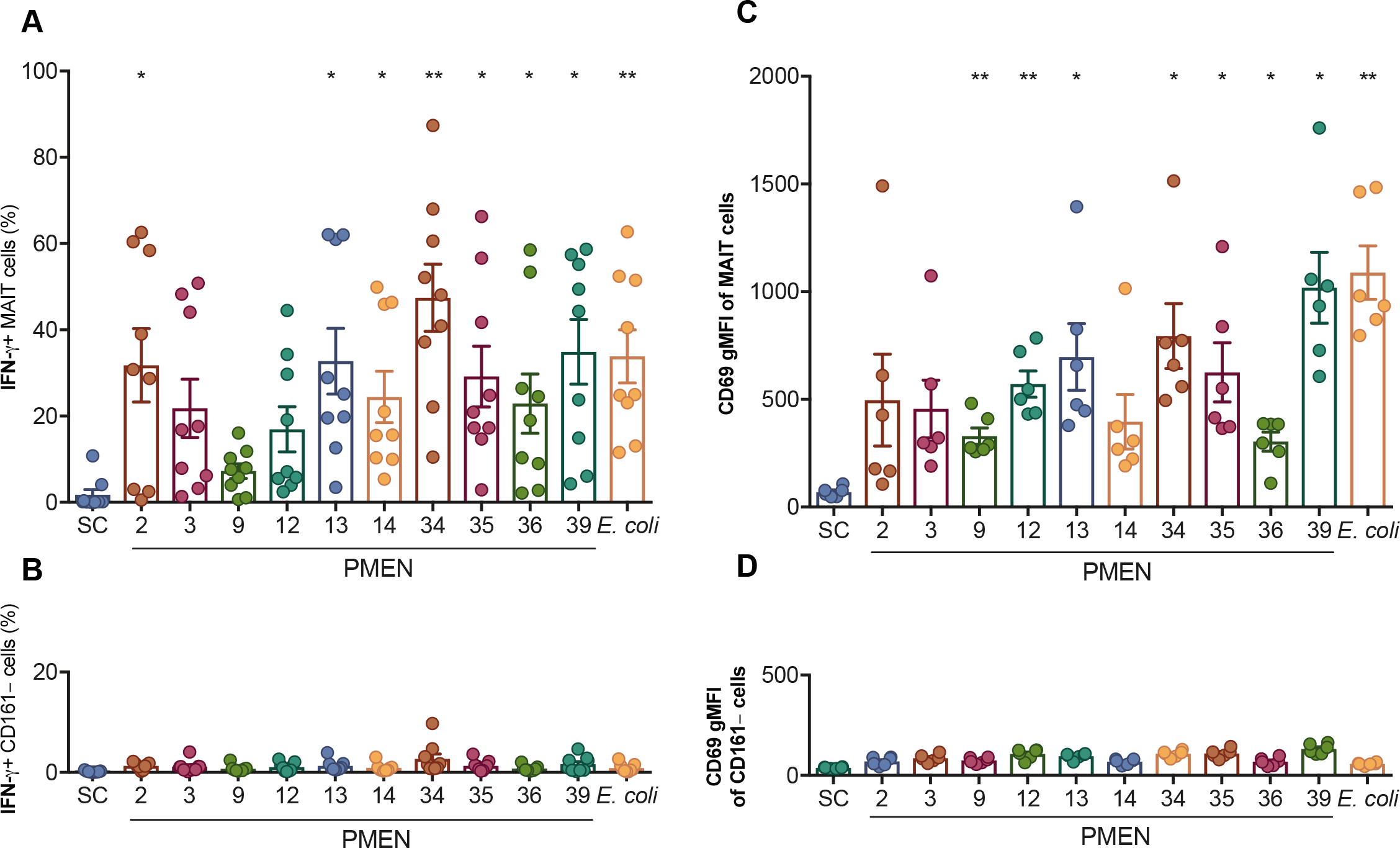
MAIT cells are activated by S. *pneumoniae*. 10 Pneumococcal Molecular Epidemiological Network (PMEN) reference strains were used to probe the activation of MAIT cells in the presence of the monocytic cell line, THP1. *Escherichia coli (E. coli)* were added as a positive control. (A-B) Frequency of cells expressing IFNγ in MAIT cells (A) or CD161-CD8+ T cells (B) are shown (n=9). (C-D) CD69 expression measured by geometric mean fluorescence intensity (gMFI) in MAIT cells (C) or CD161-CD8+ T cells (D) are shown (n=6). **P<0.01, *P<0.05 by repeated measures one-way ANOVA with Dunnett’s multiple comparisons test, compared to the sterility control (SC). Numbers indicate the PMEN reference strains. SC= sterility control.

Similarly, CD69 expression was induced by all 10 strains as measured by geometric mean fluorescence intensity (gMFI), and reached significance for 7 strains (PMEN9, PMEN12, PMEN13, PMEN34, PMEN35, PMEN36, and PMEN39). In comparison, there was negligible activation of non-MAIT cells (CD161-CD8+ T cells; Figure 2B) across all tests as measured by IFNγ or CD69 expression, suggesting that S. *pneumoniae* specifically activated MAIT cells.

### MAIT cell activation by *S. pneumoniae* in the presence of monocytes is not MR1-dependent

We have previously found that the response of MAIT cells to *E. coli* is dependent on both MR1 as well as innate cytokines IL-12 and IL-18 (14). To investigate whether the response of MAIT cells to *S. pneumoniae* was dependent on MR1, cytokines, or both, we cultured PFA-fixed S. *pneumoniae* with PBMCs and THP1 cells in the presence of anti-MR1, anti-IL-12, and anti-IL-18 blocking antibodies (Figure 3A). As expected, IFNγ expression in response to *E. coli* could be blocked significantly by anti-IL-12 and anti-IL-18 blocking antibodies, which was further reduced and abrogated by the addition of an anti-MR1 blocking antibody, showing that both the TCR and cytokine signaling pathways contribute to MAIT cell activation by *E. coli.*Surprisingly, we found that blockade of MR1 had no effect on pneumococcal MAIT cell activation. Instead, blocking IL-12 and IL-18 completely abrogated MAIT cell activation across all strains tested. This suggested that despite the fact that *S. pneumoniae* possesses the riboflavin synthesis pathway (Table 1, Figure 1A), activation of MAIT cells by *S. pneumoniae* in the presence of THP1 cells was dependent on cytokines, and not activation through antigen presentation by MR1.

**Figure 3.**
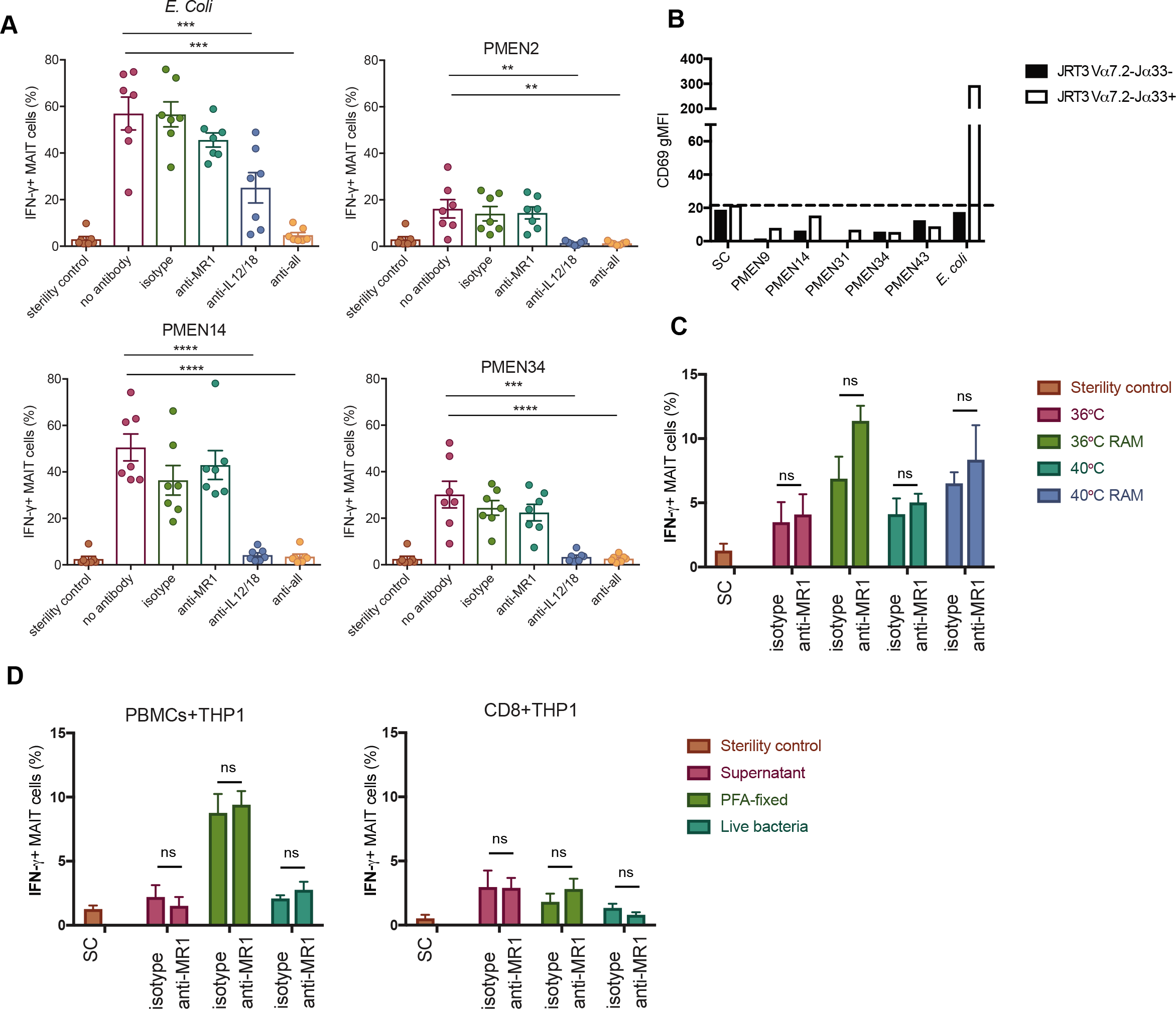
MAIT cell activation by *S. pneumoniae* in presence of monocytes is not MR1-dependent. (A) Paraformaldehyde (PFA)-fixed *S. pneumoniae* PMEN reference strains or *E. coli* were cultured with PBMCs and THP1 cells in the presence of anti-MR1, anti-IL-12, and anti-IL-18 blocking antibodies. IFNγ expression from MAIT cells are shown. ****P<0.0001, ***P<0.001, **P<0.01 by repeated measures one-way ANOVA with Dunnett’s multiple comparisons test, compared to the no antibody control (n=7). (B) Jurkat cells expressing the MAIT cell TCR (white bars) or control cells not expressing the MAIT cell TCR (black bars) were cultured with THP1 cells overnight with the indicated PMEN strains of *S. pneumoniae*, or *E. coli* as a positive control. Activation was measured as the geometric mean fluorescence intensity (gMFI) of CD69 expressed by Jurkat cells. SC=sterility control baseline activation, as indicated by the dotted line. Representative of three independent experiments. (C) The *S. pneumoniae* PMEN34 strain was grown overnight at 36°C or 40°C in THB-Y media and either cultured in THB-Y media for the last 4 hours or transferred to riboflavin-free media (RAM). The bacteria were then fixed and cultured with PBMCs and THP1 cells overnight. Frequency of IFNγ expressing MAIT cells is shown, in the presence or absence of anti-MR1 blocking antibody. ns=non-significant by two-way ANOVA with Sidak’s multiple comparisons test (n=3). (D) The *S. pneumoniae* PMEN34 strain was grown overnight and the supernatant, fixed or live bacteria were added to THP1 cells overnight with either PBMCs (left) or enriched CD8+ T cells (right), in the presence or absence of anti-MR1 blocking antibody. Frequency of IFNγ expressing MAIT cells are shown. SC=sterility control, ns=non-significant by two-way ANOVA with Sidak’s multiple comparisons test (n=3).

Next, to investigate whether *S. pneumoniae* can activate MAIT cells through the MR1 pathway alone, Jurkat cells engineered to express the MAIT cell TCR Vα7.2-Jα33 were cultured with fixed *S. pneumoniae* strains and cultured overnight in the presence of THP1 cells (Figure 3B). There was no significant change in the expression of CD69 by MAIT-Jurkats in the presence of any of the *S. pneumoniae* strains, although the cells were able to respond to the *E. coli* positive control. Thus, in the presence of monocytic cells, there was very little activation of MAIT cells by *S. pneumoniae* through the MR1 pathway.

Given the upregulation of the riboflavin synthesis pathway in *S. pneumoniae* upon heat stress at 40°C (Figure 1B), we tested whether changing environmental factors such as temperature and availability of riboflavin in the growth media would trigger activation of MAIT cells through the MR1 pathway. *S. pneumoniae* strain PMEN34 was grown for 16 hours in Todd Hewitt Broth (THB) media supplemented with yeast (THB-Y) at 36°C and then either transferred to 40°C incubation, or transferred to riboflavin-free assay media (RAM) at 36°C or 40°C for 4 hours, before the bacteria were fixed (Figure 3C). Although there was a slight increase in the fraction of MAIT cells expressing IFNγ when bacteria were cultured in the absence of riboflavin in RAM, regardless of temperature, this increase was not dependent on MR1 as the addition of an anti-MR1 blocking antibody did not reduce the frequency of IFNγ-expressing MAIT cells. Thus, the complete absence of riboflavin, increased incubation temperature or the combination of both did not lead to MR1-dependent activation of MAIT cells.

We also tested whether using live *S. pneumoniae* strain PMEN34 or the supernatant of *S. pneumoniae* growth culture, instead of fixed bacteria, would stimulate MAIT cells through the MR1-pathway (Figure 3D); however, these responses were small and could not be significantly blocked by an anti-MR1 blocking antibody. We enriched for CD8+ T cells and added these to the assay instead of PBMCs in case other cells were interfering with MAIT cell activation through MR1, but these responses were similar to those using PBMCs and not affected by MR1-blocking. Thus, in the presence of monocytes, MAIT cells are activated mainly through innate cytokines rather than MR1, regardless of temperature or riboflavin availability.

### MR1-dependent activation of MAIT cells by *S. pneumoniae* in the presence of macrophages

We next tested whether monocyte-derived macrophages may be able to present the MRI-ligand to activate MAIT cells more effectively through MR1, as alveolar macrophages play an important role in the immune response to *S. pneumoniae*(21,22). Furthermore, we investigated whether temperature or the abundance of riboflavin in the media affects MR1-dependent activation of MAIT cells in the presence of macrophages. For this, *S. pneumoniae* strain PMEN34 was grown for 16 hours in THB media, with or without yeast extract (THB-Y and THB, respectively) at either 36°C or 40°C. Given that the riboflavin synthesis pathway was upregulated by a short incubation at 40°C (Figure 1), we also transferred half of the bacteria grown overnight at 36°C to 40°C for 4 hours. The bacteria were fixed immediately and added to PBMCs and monocyte-derived macrophages overnight (Figure 4A,B).

**Figure 4.**
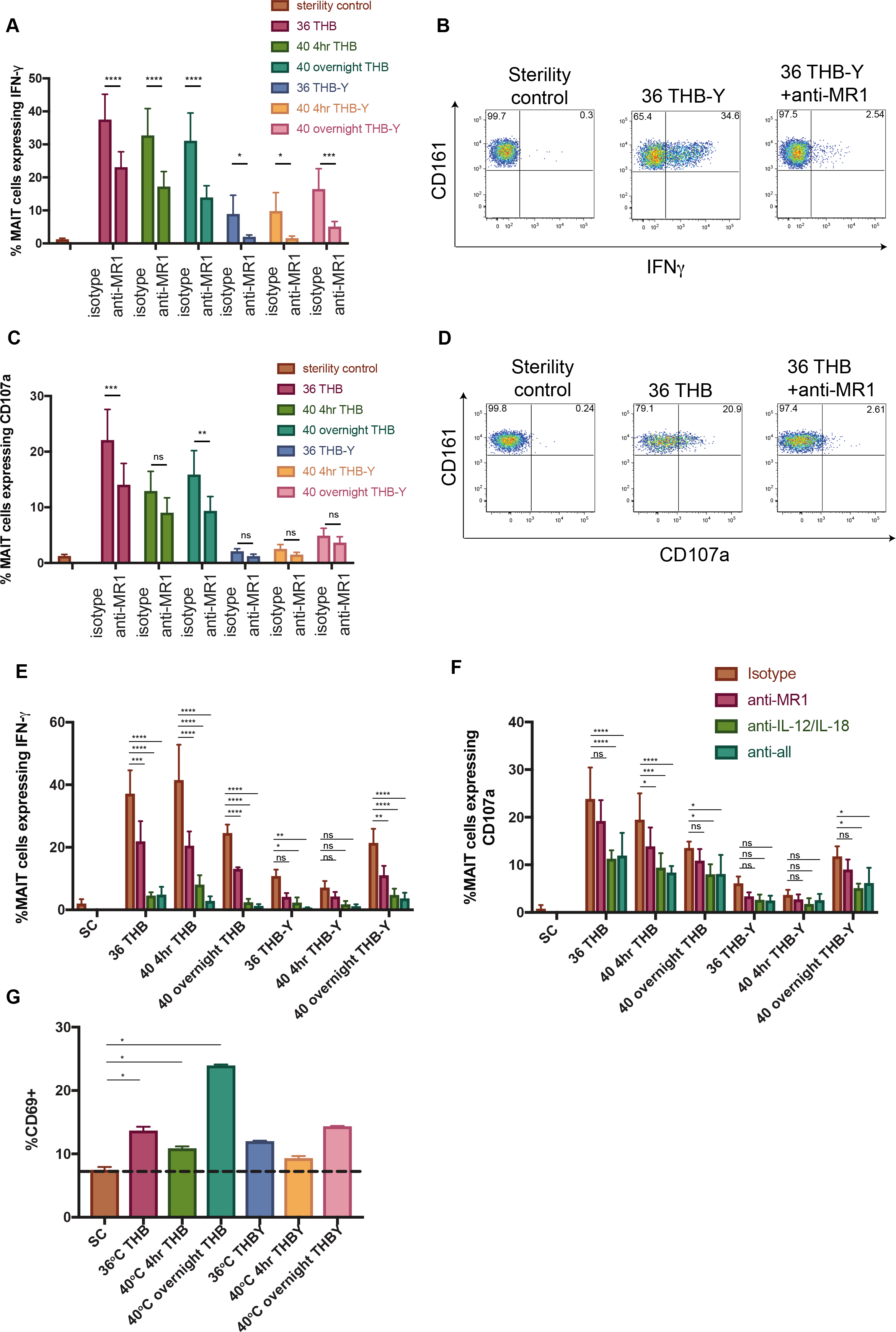
MR1-dependent activation of MAIT cells by *S. pneumoniae* in the presence of macrophages. A-D) The *S. pneumoniae* PMEN34 strain was grown for 16 hours in THB media, with or without yeast extract (THB-Y and THB, respectively) at either 36°C (36) or 40°C (40 overnight). Half of the bacteria grown overnight at 36°C were transferred to 40°C for 4 hours (40 4 hr). The bacteria were fixed immediately and added to PBMCs and monocyte-derived macrophages overnight, in the presence or absence of anti-MR1 blocking antibody. Frequency of MAIT cells expressing IFNγ (A) and representative example of IFNγ expression from MAIT cells by FACS (B) are shown with isotype control or anti-MR1 blocking antibody. ****P<0.0001, ***P<0.001, and * P<0.05 by two-way ANOVA with Sidak’s multiple comparisons test (n=6). Frequency of MAIT cells expressing CD107a (C) and representative example of CD107a expression from MAIT cells by FACS (D) are shown with isotype control or anti-MR1 blocking antibody. ***P<0.001, **P<0.01, or non-significant (ns) by two-way ANOVA with Sidak’s multiple comparisons test (n=4). 36=36°C, 40=40°C. E-F) The *S. pneumoniae* PMEN34 strain was grown for 16 hours in THB-Y or THB media, at either 36°C or 40°C. Half of the bacteria grown overnight at 36°C were transferred to 40°C for 4 hours. The bacteria were fixed immediately and added to PBMCs and monocyte-derived macrophages overnight, in the presence or absence of indicated combinations of anti-MR1, anti-IL-12, and anti-IL-18 blocking antibodies. Frequency of MAIT cells expressing IFNγ (E) and CD107a (F) are shown. 36=36°C, 40=40°C. ****P<0.0001, ***P<0.001, **P<0.01, *P<0.05 or non-significant (ns) by two-way ANOVA with Dunnett’s multiple comparisons test (n=3). (G) Jurkat cells expressing the MAIT cell TCR were cultured with monocyte-derived macrophages overnight with the PMEN34 strain of *S. pneumoniae.* Activation was measured as frequency of Jurkat cells expressing CD69. Dotted line indicates CD69 expression by Jurkats in the presence of sterility control (SC). THB=Todd Hewitt Broth, THB-Y=THB with yeast extract. *P<0.05 by repeated measures one-way ANOVA with Dunnett’s multiple comparisons test. All experiments were performed in duplicates, and representative of two independent experiments.

Surprisingly, we found that when using monocyte-derived macrophages, *S. pneumoniae* induced IFNγ expression from MAIT cells that was reduced by an anti-MR1 blocking antibody, suggesting that the response was MR1-dependent, in contrast to the response in the presence of monocytes. The addition of the anti-MR1 blocking antibody significantly reduced the frequency of IFNγ-expressing MAIT cells regardless of the temperature and media in which the *S. pneumoniae* were grown. Interestingly, there was a clear increase in activation induced by bacteria grown in THB media without the addition of yeast, compared to bacteria grown in the presence of yeast. Degranulation was also induced by *S. pneumoniae* grown in THB media and was blocked by the anti-MRI-blocking antibody to a varying degree (Figure 4C,D).

Next, in order to compare the dependency of MAIT cell activation by *S. pneumoniae*on MR1, IL-12 and IL-18 in the context of macrophages, *S. pneumoniae* strain PMEN34 was grown for 16 hours in THB or THB-Y media at either 36°C or 40°C. Half of the bacteria grown overnight at 36°C were transferred to 40°C for an additional 4 hours. Then the bacteria were fixed and added to PBMCs and macrophages overnight in the presence of anti-MR1, anti-IL-12, and anti-IL-18 antibodies (Figure 4E, F). We found that there was a significant effect of blocking MR1 on IFNγ production from MAIT cells in the presence of *S. pneumoniae* cultured in THB media, regardless of temperature, but this was almost completely abrogated in the presence of anti-IL-12 and anti-IL-18 blocking antibodies. There was a similar effect of blocking antibodies on MAIT cell IFNγ production in response to *S. pneumoniae* cultured in THB-Y media overnight at 40°C. Furthermore, although we have previously found that degranulation in response to *E. coli*was completely dependent on MR1 and not on cytokines (23), degranulation of MAIT cells in response to *S. pneumoniae* was, in each case, further reduced by the addition of anti-IL-12 and anti-IL-18 compared to blocking MR1 alone. In particular, degranulation in response to *S. pneumoniae* grown at 40°C for the last 4 hours in THB media was significantly reduced by anti-MR1 blocking, which was further reduced by the addition of anti-IL-12 and anti-IL-18 blocking antibodies (Figure 4F). This suggests that degranulation in response to *S. pneumoniae* requires co-stimulation from cytokines.

Finally, we set out to confirm that *S. pneumoniae* were able to activate MAIT cells in the presence of macrophages in an MRI-dependent manner. We cultured Jurkat cells that express the MAIT cell TCR Va7.2-Jα33 with fixed *S. pneumoniae* grown in THB or THB-Y media at different temperatures in the presence of monocyte-derived macrophages (Figure 4G). This showed that *S. pneumoniae* grown in THB media significantly increased the expression of CD69 in MAIT-Jurkat cell lines, compared to the Jurkat cells cultured with the sterility control, although the response was relatively weak. Thus, in the presence of monocyte-derived macrophages, *S. pneumoniae* were able to activate MAIT cells in an MRI-dependent manner, but were highly dependent on the presence of cytokines to boost the response.

### Riboflavin operons are also present in non-pneumococcal *Streptococcus* spp

A bioinformatic investigation of 824 genomes of 69 different *Streptococcus* spp. revealed that the riboflavin operon was also present in other streptococci. 11 different versions of the riboflavin operon were identified among 13 non-pneumococcal *Streptococcus* spp. (Supplementary Table 2). The majority of these riboflavin operons were located between genes involved in arginine biosynthesis (*argC, argJ, argB, argD)* and ribonucleotide reduction *(nrdF2, nrdE2, nrdH)* (Figure 5A). Despite identical gene synteny between different versions of the riboflavin operon (Figure 5A), they differed greatly in nucleotide sequence identity (Figure 5B).

**Figure 5.**
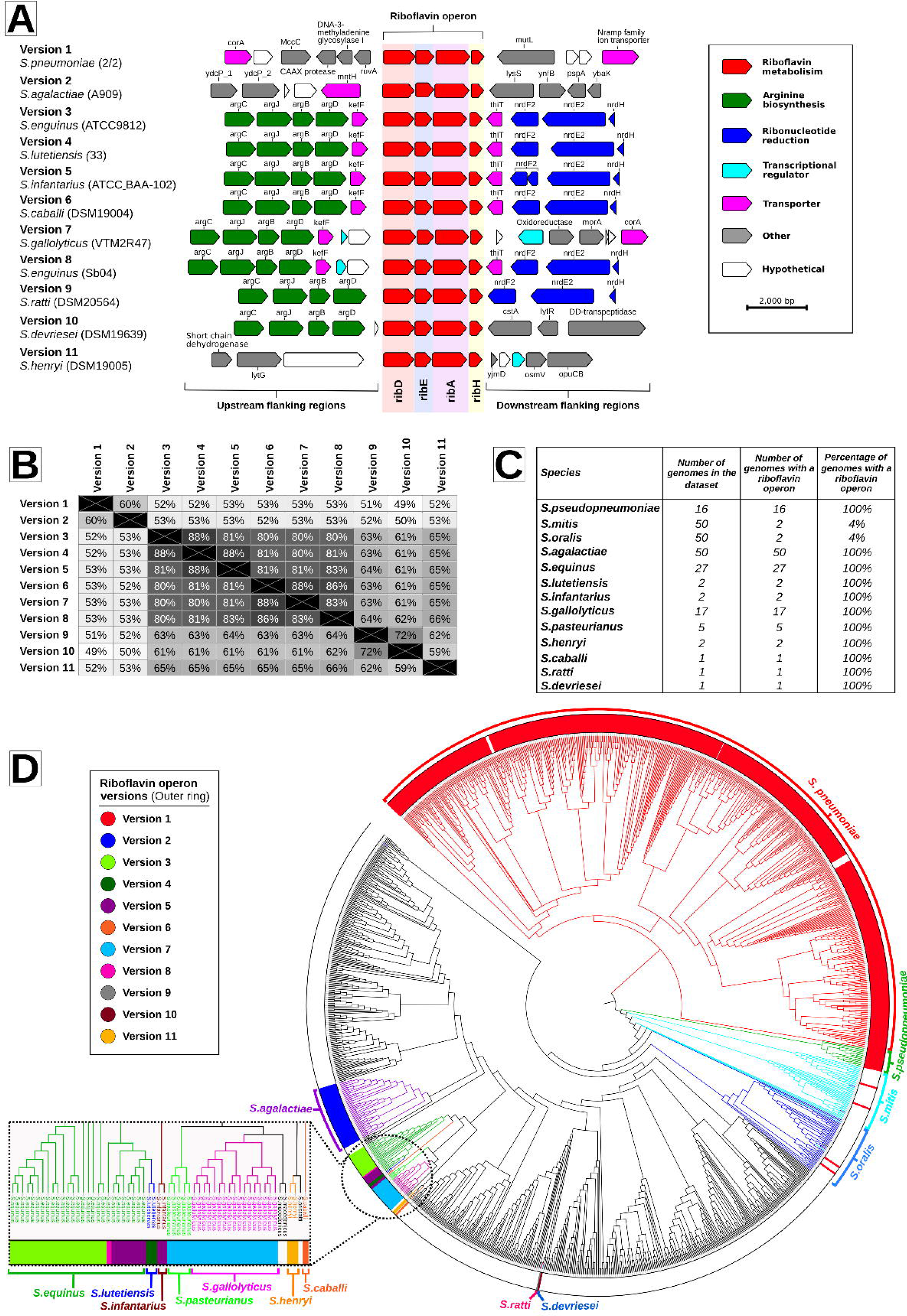
Evidence for different versions of riboflavin operons in other *Streptococcus* spp. A) The riboflavin operon found in *S. pneumoniae* (version 1) and its flanking genes are depicted and compared to 10 additional representative versions of the riboflavin operon found among other *Streptococcus* spp. B) Matrix of pairwise comparisons of nucleotide similarity among the 11 different versions of the riboflavin operon. C) Table summarizing the riboflavin operons found in 13 non-pneumococcal *Streptococcus* spp. D) Phylogenetic tree constructed based upon the concatenated sequences of 53 ribosomal MLST loci among 571 *S. pneumoniae* and 824 *Streptococcus* spp. genomes. Branches of the tree were coloured grey if no riboflavin operon was identified within the genome, whereas other colours represent genomes in bacterial species that did possess a version of a riboflavin operon. The coloured outer ring indicates the version of riboflavin operon that was identified in each genome or set of genomes. The rectangular box contains an expanded view of the circled area of the phylogenetic tree.

The *S. pneumoniae* version of the riboflavin operon was found among all 16 genomes of S. *pseudopneumoniae* and two genomes each of S. *mitis* and S. *oralis*(Figure 5C-D). No riboflavin operon genes were identified among the remaining 48 S. *mitis* and 48 S. *oralis* genomes. S. *pneumoniae, S pseudopneumoniae, S. mitis* and S. *oralis* are all closely-related commensal streptococcal species that can exchange DNA with one another; therefore the limited number of riboflavin operons present in *S. mitis* and *S. oralis* suggest that the examples identified here were the result of horizontal genetic exchange (24). Some versions of riboflavin operons identified among other *Streptococcus* spp. were exclusively present in only one species, e.g. version 2 was identified in all 50 genomes of S. *agalactiae* but no other species; whereas in contrast, S. *equinus* contained three different versions of the riboflavin operon, one of which (version 5) was also found in S. *infantarius* (Figure 5D; Supplementary Table 2). Furthermore, over 2400 S. *pyogenes* genomes were investigated for the presence of Rib genes. The riboflavin operon could not be found in the genomes of S. *pyogenes* (data not shown).

## Discussion

Our study is the first to demonstrate that MAIT cells can respond to and recognize S. *pneumoniae.* Given the urgent global need to tackle antimicrobial-resistant strains and pneumococci not covered by the currently available vaccines, understanding the mechanism by which MAIT cells are activated by *S. pneumoniae* provides a method of targeting a metabolic pathway that is highly conserved amongst S. *pneumoniae.*

We found that MAIT cell activation by *S. pneumoniae* through the MR1-restricted pathway was dependent on the type of antigen-presenting cell. There was a significant effect of blocking MR1 recognition by MAIT cells only when the *S. pneumoniae* was presented by macrophages, but not monocytes (either those amongst PBMCs or by the monocytic cell line THP1). Indeed, the lack of MAIT cell activation by *S. pneumoniae* in a previous paper (25) is in line with our finding that monocytes are not able to sufficiently activate MAIT cells solely through the presentation of ligands from *S. pneumoniae* by MR1. The reason for the difference in the ability of these antigen-presenting cells to activate MAIT cells through MR1 antigen presentation in the case of *S. pneumoniae* is not clear. It has been suggested that monocytes are poor antigen presenters (26) and instead function as antigen-transporting cells that deliver antigen to the draining lymph nodes (27). Although THP1 cells and monocytes have been used repeatedly to activate MAIT cells in an MR1-dependent manner with numerous other bacteria, the pathogenesis of *S. pneumoniae* is related to its many virulence factors, some of which are associated with minimizing phagocytosis and thereby antigen presentation (further discussed below). Macrophages may be a more powerful antigen-presenting cell that are able to efficiently phagocytose the bacteria, provide more co-stimulatory signals, or resist interference with antigen presentation and loading of MR1. Indeed, mice lacking alveolar macrophages, located at the interphase between air and lung tissue, show a delayed and impaired bacterial clearance as compared with control mice (21,22). Thus, alveolar macrophages provide the first line of cellular defense against microbes and may be an important presenter of *S. pneumoniae* antigen to MAIT cells.

Even in the presence of macrophages, the MAIT cell response to *S. pneumoniae*was highly co-dependent on cytokines. This is most likely due to the weak TCR signal induced by *S. pneumoniae*, as seen by the significant but weak activation of MAIT-Jurkat cell lines even when using macrophages. MAIT cells are characterized by downregulated TCR signaling components and a high dependency on co-stimulation provided by cytokines for activation and expansion (28–30). The particularly weak TCR signal provided by *S. pneumoniae* may be because *S. pneumoniae* is a bacterium highly specialized to evade the host immune system by circumventing phagocytosis and antigen presentation. The efficiency of phagocytosis can be impeded by the presence of the thick polysaccharide capsule, while another virulence-associated trait is the ability of this bacterium to undergo autolysis. Autolysis of *S. pneumoniae* has been shown to specifically reduce the phagocytosis of intact bacteria, even in the absence of a polysaccharide capsule, and reduce the production of phagocyte-activating cytokines such as TNFα, IFNγ and IL-12 (31). The uniquely high sensitivity of MAIT cells to cytokines such as IL-12 (32) allows these cells to boost the immune response and provide early IFNγ production.

The biggest disease burden caused by *S. pneumoniae* by far is pneumonia, which is the leading infectious cause of mortality in children under five years of age (1), as well as accounting for a large part of community-acquired pneumonia in the elderly (2). Given that we show the mechanism by which MAIT cells respond to *S. pneumoniae*, and the critical role that pulmonary MAIT cells play *in vivo* against lung infections (18,20), it would be reasonable to suggest that MAIT cells may play a role in pneumococcal pneumonia. These cells may also be a critical factor in secondary bacterial pneumonia that follow influenza infection, which are most commonly caused by *S. pneumoniae* and associated with high mortality (33). MAIT cells are reduced in frequency in the blood of patients with acute influenza infection, particularly in those who succumbed to this disease (15,34). The loss of MAIT cells due to primary influenza infection may leave the patient vulnerable to secondary infection by *S. pneumoniae*, due to the lack of early control and immunomodulation provided by these cells, as has been shown with γδ T cells (35). Whether the low numbers of MAIT cells in neonates (36,37), or the decline in MAIT cell numbers in the elderly (38) and in influenza (15) affects the susceptibility of these patients to pneumococcal pneumonia will be important to investigate in *in vivo* models of pneumococcal infection (39).

Interestingly, we found that temperature and riboflavin availability for *S. pneumoniae*have the potential to affect riboflavin synthesis and MAIT cell activation. In our models using macrophages, there was a difference in MR1-dependent MAIT cell activation induced by *S. pneumoniae* cultured in THB media compared to THB-Y media, the latter of which is supplemented with riboflavin-rich yeast extract. Although the THB media alone supports the growth of riboflavin-auxotrophs such as *Streptococcus pyogenes*, this has been shown to be due to the ability of S. *pyogenes* to utilize THB-derived components as a substitute for riboflavin, and mutant strains without this ability require the addition of riboflavin or riboflavin-rich yeast in order to grow in THB media (40). Thus, THB media on its own does not contain riboflavin. We found that *S. pneumoniae* grown in THB media consistently induced greater MAIT cell activation, which may be partly due to the increased production of riboflavin in bacteria grown in riboflavin-deficient THB media, compared to those grown in riboflavin-rich THB-Y media. We also found that riboflavin synthesis among *S. pneumoniae* was influenced by temperature: RNA-seq transcriptomic analyses of *S. pneumoniae* revealed that incubation at 40°C upregulated the riboflavin synthesis genes. A relationship between riboflavin production and temperature has been demonstrated before in *Lactococcus lactis:* it suffers from oxidative stress caused by riboflavin starvation at high temperatures and results in severe growth inhibition, which can be improved by the addition of riboflavin to the medium (41). Although no consistent increase in MR1-dependent MAIT cell activation could be detected in response to pneumococci grown at higher temperatures in our short-term activation models, it may be possible that MAIT cells are able to react to increased riboflavin production of *S. pneumoniae* at higher body temperatures during a fever. Thus, MAIT cells may be able to not only recognize the presence of *S. pneumoniae* but respond to the environment and condition in which the bacteria is growing - an important indicator of the pathogenic potential of a commensal pathogen such as *S. pneumoniae.*

We used a population genomics approach to assess the prevalence and diversity of the riboflavin operon among a large and diverse collection of S. *pneumoniae.* This revealed that the riboflavin genes are nearly ubiquitous and highly conserved at a nucleotide level among *S. pneumoniae* that had been recovered over the past century. This provides confidence that the findings presented here are generalizable among *S. pneumoniae*, with rare exceptions.

We also assessed the presence of riboflavin synthesis genes among non-pneumococcal *Streptococcus* spp. and found that a number of other species do possess these genes, including other human-associated commensal streptococci like *S. agalactiae.* Notably though, not all *Streptococcus* spp. possessed a riboflavin operon (at least one that was detectable by our screening method) and there was variation among the genomes that did, either in that more than one version of riboflavin operon was detected within a bacterial species, or that the presence of a riboflavin operon was not ubiquitous among genomes of a particular bacterial species. Hence, caution must be exercised when extrapolating findings based on a small number of bacterial strains to the population as a whole since they may not be representative, and ideally a population-based approach is used whenever possible. However, the fact that other *Streptococcus* spp. also possess a riboflavin operon presents opportunities for future studies. For example, given that MAIT cells reside in the female genital mucosa (42) it would be important to explore whether there is a MAIT cell response in the context of vaginal colonization of S. *agalactiae* (group B streptococci) among pregnant women and invasive neonatal infections (43).

Overall these data show a robust response of MAIT cells to *S. pneumoniae*, and conservation of the relevant biosynthetic pathway in this organism and other closely-related *Streptococcus* spp. Given the low levels of MAIT cells in early life and their decline in old age – the highest risk populations for invasive pneumococcal disease – further understanding of the functional role of MAIT cells *in vivo* in host defense against this major pathogen should be of substantial interest.

## Materials and Methods

### Flow cytometry

For immunofluorescence staining, dead cells were excluded with the Live/Dead Fixable near-IR dead-cell stain (Invitrogen). For internal staining, cells were fixed with 1% formaldehyde (Sigma Aldrich) and permeabilised with permeabilisation buffer (eBioscience). Antibodies used were: CD3 Pacific Orange, Granzyme B APC (Life Technologies), CD8 PE-Cy7, TNFα PerCP-Cy5.5, Vα7.2 APC, FITC, or PE, CD107a PE-Cy7 (BioLegend), CD3 efluor 450, CD8 PerCP-Cy5.5, CD69 FITC or Pacific Blue, IFNγ PerCP-Cy5.5 (eBiosciences), CD161 PE, CD4 VioGreen, IFNγ FITC (Miltenyi Biotec).

### Cells

Whole blood was obtained from leukocyte cones (NHS Blood and Transplant). PBMCs were isolated by standard density gradient centrifugation (Lymphoprep™ Axis Shield Diagnostics). All samples were collected with appropriate patient consent and local research ethics committee approval (COREC 04.OXA.010). Where indicated, CD8+ cells were positively enriched using CD8 Microbeads (Miltenyi Biotech) before *in vitro* stimulation. Monocyte-derived macrophages were generated by enriching for monocytes using CD14 Microbeads (Miltenyi Biotech) before culturing with 50ng/ml GM-CSF (Miltenyi Biotech) in RPMI media, penicillin/streptomycin, L-glutamine, and 10% human serum for 6-8 days.

The Jurkat-MAIT cell line was produced by cloning into a pHR-IRES vector the TRAV1-2-TRAJ33 (CDR3α: CAVMDSNYQLIW) MAIT cell α chain and then cloning in the TRBV6-1-TRBJ2-3 β chain (CDR3β-CASSETSGSPDTQYF). The vector was then co-transfected into 293T cells with the HIV gag-pol and VSV-G expression plasmids using X-tremeGENE™ 9 DNA Transfection Reagent (Sigma) according to the manufacturer’s instructions. The supernatant from this culture containing the lentiviral particles was then used to transduce J.RT3-T3.5 (JRT3) cells (which lack an endogenous TCRβ chain). TCR expression and pairing with MAIT cell TRAV1-2 (Va7.2) chain was confirmed by flow cytometry and cells sorted based on CD3 and Va7.2 expression.

### Pneumococcal reference strains used in experiments

Ten PMEN reference strains of *S. pneumoniae* were tested in this study (note: country of first detection^serotype^; multilocus sequence type): PMEN2 (Spain^6B^-2; ST90), PMEN3 (Spain^9V^-3; ST156), PMEN9 (England^14^-9; ST9), PMEN12 (Finland^6B^-12; ST270), PMEN13 (S.Africa^19A^-13; ST41), PMEN14 (Taiwan^19F^-14; ST236), PMEN34 (Denmark^12F^-34; ST218), PMEN35 (Netherlands^14^-35; ST124), PMEN36 (Netherlands^18C^-36; ST113), PMEN39 (Netherlands^7F^-39; ST191)(44–46). Eight serotypes were represented, all of which are vaccine serotypes apart from 12F (PMEN34).

PMEN strains were cultured from freezer stocks to Columbia blood agar plates (Oxoid) and incubated overnight at 37°C+5% CO_2_, then transferred to THB media (Sigma Aldrich) with 0.5% yeast extract (THB-Y; Sigma Aldrich) and incubated at 37°C + 5% CO_2_ overnight, unless indicated otherwise. In specific experiments, bacteria were grown in Riboflavin Assay Media (RAM; BD Difco). *E. coli*(DH5a, Invitrogen) were cultured in Lysogeny Broth (LB) medium and incubated overnight in a shaking incubator.

*S. pneumoniae* or *E. coli* were fixed in 1% PFA for 15 minutes, and washed extensively in sterile, filtered phosphate-buffered saline (PBS). A negative control was prepared in identical fashion and used as a sterility control. Following fixing, bacteria were immediately used in *in vitro* stimulation assays. Alternatively, filtered bacterial culture supernatant or unfixed live bacteria was added to assays where indicated. Live pneumococci were added to PBMCs and antigen-presenting cells for 30 minutes at 37°C before cells were extensively washed and cultured for the remainder of the overnight assay in 100μg/ml gentamycin.

### *In vitro* stimulation of MAIT cells

THP1 cells (ECACC, UK) were incubated overnight with PFA-fixed *S. pneumoniae* or *E. coli* at a ratio of 30 bacteria per cell (BpC), or the sterility control (without bacteria, see above). THP1 cells were washed and PBMCs or enriched CD8+ T cells were added to THP1 cells overnight. Brefeldin A (eBioscience) was added for the final 4 hours of the stimulation. Alternatively, for the assessment of degranulation, anti-CD107a PE-Cy7 (BioLegend) was added from the start of the stimulation. For blocking experiments, anti-MR1 (BioLegend), anti-IL-12p40/70, anti-IL-18 antibody (both BioLegend), or the appropriate isotype controls, were added for the duration of the experiment.

### Data acquisition and statistical analysis

Data on MAIT cell activation were collected on the MACSQuant Analyser (Miltenyi Biotech) and were analysed using FlowJo v9.8 (TreeStar). All graphs and statistical analyses were completed using GraphPad Prism software version 6. All data are presented as means with standard error of the mean (S.E.M.) unless otherwise indicated.

### Compilation of the genome datasets

Two large genome datasets were compiled for this study and data were stored in a BIGSdb database (47). The *S. pneumoniae* dataset consisted of 571 historical and modern genomes isolated from 1916-2009 from people of all ages residing in 39 different countries. The *S. pneumoniae* isolates were recovered from both carriage and disease, and 89 serotypes and 296 multilocus sequence types were represented in this dataset (Supplementary Table 1). 486 *S. pneumoniae* genome sequences were compiled from previously published studies or were downloaded from GenBank (48). The remaining 85 *S. pneumoniae* genomes were recently sequenced. *S. pneumoniae* cultures were prepared and incubated as described above, before DNA was extracted using the Promega Maxwell^®^ 16 Instrument and Buccal Swab LEV DNA purification kits following the manufacturer’s protocol. DNA extracts were sent to the Oxford Genomics Centre where libraries were made and DNA was sequenced on the Ilumina^®^ platform. Velvet was used to make *de novo* genome assemblies, which were further improved using SSPACE and GapFiller (49–51).

The non-pneumococcal *Streptococcus* spp. dataset contained 834 genomes of 69 different streptococcal species (Supplementary Table 2). 34 genomes were newly sequenced as above and the rest were downloaded from the ribosomal multilocus sequence typing (rMLST) database (52). For each species, the number of genomes included in this study dataset was capped at 50: if fewer than 50 genomes were available in the rMLST database for a given species, then all available genomes were included, but if more than 50 sequenced genomes were available then genomes were manually selected for inclusion. In these instances, the population structure of the species was depicted using PHYLOViZ and 50 genomes were selected with the aim of maximising the population-level diversity of that species from the available genomes (53).

Genomes were annotated using both RAST (54) and Prokka (55). Genome sequences can be accessed from the European Nucleotide Archive (56), PubMLST website(57), GenBank(58) and/or the rMLST database(59) (see Supplementary Tables 1 and 2).

### RNA-sequencing experiments and analyses

*S. pneumoniae* isolate 2/2 was cultured in seven 10 ml tubes of brain-heart infusion broth and incubated at 40°C + 5% CO_2_ for 6 hours to mimic heat shock. The experimental control was the same isolate 2/2, also cultured in seven 10 ml tubes of brain-heart infusion broth but incubated at standard conditions of 37°C + 5% CO_2_. Broth cultures at five time points (2, 3, 4, 5 and 6 hours of incubation) were removed and 19 ml of RNAprotect Bacteria Reagent (Qiagen) was added to stabilise the RNA. RNA was extracted from the samples using the Promega Maxwell^®^ 16 Instrument and LEV simplyRNA Cells purification kit, following the manufacturer’s protocol. Extracted RNA samples were sent to the Oxford Genomics Centre for processing. Library preps were made using RNA-Seq Ribozero kits (Illumina, Inc) and sequencing was performed on the MiSeq (Illumina, Inc).

The sequenced forward and reverse reads were paired and mapped to the *S. pneumoniae* strain 2/2 genome using Bowtie2 with the highest sensitivity option (60). Differential gene expression was analysed in Geneious version 9.1 (Biomatters Ltd) using the DESeq method (61). Genes with an adjusted p-value < 0.05 were considered to be differentially expressed. RNA sequencing data used in this paper have been deposited in the Gene Expression Omnibus (GEO) database with accession number GSMXXXXXXX (pending).

### Genomic analyses of riboflavin operon genes

Genes involved in riboflavin metabolism were identified using the KEGG pathway database (KEGG entry number: snp00740) and previously published experimental work (12,62). Individual BLAST searches of *ribD, ribE, ribA* and *ribH* sequences among all 571 *S. pneumoniae* genomes were performed via the BIGSdb database. There were two instances each for *ribD* and *ribH* where the gene sequences were split over multiple assembly contigs and these sequences were excluded from further analyses. Individually, multiple nucleotide sequence alignments for *ribD, ribE, ribA* and *ribH* were performed in Geneious using the ClustalW algorithm with default parameters (Gap open cost=15, Gap extend cost=6.66) (63). To compute the dN/dS ratio, the number of synonymous (dS) and non-synonymous (dN) substitutions per site was determined on codon-aligned sequences using a maximum likelihood method, conducted through HyPhy (64) on the Datamonkey server (65,66). For the pairwise positive selection analysis, amino acid sequences were aligned in Geneious using the ClustalW algorithm (Cost matrix: BLOSUM, Gap open cost=10, Gap extend cost=0.1).

### Identification and categorisation of the riboflavin operon genes in non-pneumococcal *Streptococcus* spp

The *ribD, ribE, ribA* and *ribH* sequences in *S. pneumoniae* strain 2/2 genome were used as the query to BLAST against the non-pneumococcal *Streptococcus* spp. genome dataset (parameters: word size 11; reward 5; penalty −4; gap open 8; gap extend 6). All BLAST hits were manually inspected to confirm the presence of the riboflavin genes. Protein domains in the identified genes were annotated using the Conserved Domain search feature at NCBI (67) to confirm the presence of riboflavin biosynthesis genes. To categorise the different versions of the riboflavin operon, all sequences were clustered using CD-Hit (68) at a ≥90% similarity threshold, and a representative sequence from each cluster was selected. One ‘cluster’ contained a single riboflavin operon sequence that was disrupted by a transposon and this single sequence was excluded from further analyses. Multiple nucleotide sequence alignments of different versions of the riboflavin operons was performed in Geneious using the ClustalW algorithm with default parameters (Gap open cost=15, Gap extend cost=6.66). The multiple nucleotide sequence alignment output was used within the Geneious environment to calculate the percentage identity matrix.

### Assessment of the relationships among *Streptococcus* spp

A phylogenetic tree was built using concatenated sequence data from the ribosomal loci using the neighbour-joining method as implemented using the BIGSdb PhyloTree plugin (52,69). The tree was annotated using iTOL (70) and Inkscape (71).

## Acknowledgements

The authors gratefully acknowledge Professor Regine Hakenbeck at the University of Kaiserslautern for the stock cultures of streptococci that were newly sequenced in this study.

## Supporting information captions

Supplementary Table 1. Descriptive data for the 571 *Streptococcus pneumoniae* genomes included in this study.

Supplementary Table 2. Descriptive data for the 824 non-pneumococcal *Streptococcus* species genomes included in this study.

